# BEHAV3D-Explorer enables natural language-assisted profiling of 3D cellular dynamics

**DOI:** 10.64898/2026.11.06.764820

**Authors:** Sam de Blank, Emilio Rios-Jimenez, Alejandra Rubio-Muñoz, Oliver Hoffman, Britt M. T. Vervoort, Amber K.L. Wezenaar, Jessica K. Mazalo, Miguel Molina-Moreno, Hsin-Fang Chang, Ellen J Wehrens, Anne C. Rios, Maria Alieva

## Abstract

Obtaining quantitative measurements from live 3D microscopy remains difficult to implement across research settings, particularly without profound image-analysis expertise. We present BEHAV3D-Explorer, an interactive, user-friendly platform that uses natural-language guidance to help users select analysis strategies, tune parameters and configure workflows. BEHAV3D-Explorer enables time-resolved behavioral classification and integrates it with morphological, interaction and functional analyses, thereby connecting dynamic readouts to yield interpretable results across imaging applications.

## Main

Live 3D microscopy enables the investigation of cellular dynamics across experimental systems, from *in vitro* organoid models and tissue explants to living animals, and is increasingly used to advance therapeutic development and evaluate treatment responses^1^. However, extracting quantitative insights from volumetric time-lapse datasets remains challenging. Technical differences in imaging modalities and spatial and temporal resolution, together with heterogeneity in cellular composition, morphology and dynamics within and across biological systems, demand adaptable segmentation, tracking and analysis workflows^2–8^. Selecting, configuring and fine-tuning these workflows to individual datasets remains a major barrier to their routine use in biological research. Moreover, dynamic cellular features captured by live imaging are often analyzed independently, limiting their integration into a coherent understanding of cellular behavior and function.

Recent advances in language-model-assisted bioimage analysis have demonstrated the potential of natural-language interfaces for image interpretation and analysis workflow generation^9,10^. Here, we introduce BEHAV3D-Explorer, an open-source, end-to-end platform dedicated to live 3D microscopy that combines modular image-analysis workflows, interactive quality control and natural-language guidance through QueenB in a code-free graphical user interface (GUI) (**Fig. 1a**, **Supplementary Table 1**). Designed to make advanced quantitative analysis accessible without extensive computational expertise, BEHAV3D-Explorer extends the previously described BEHAV3D framework^2,3^, by enabling time-resolved quantification of cellular behavioral states and their integration with morphological, interaction and functional readouts.

**Figure 1.**
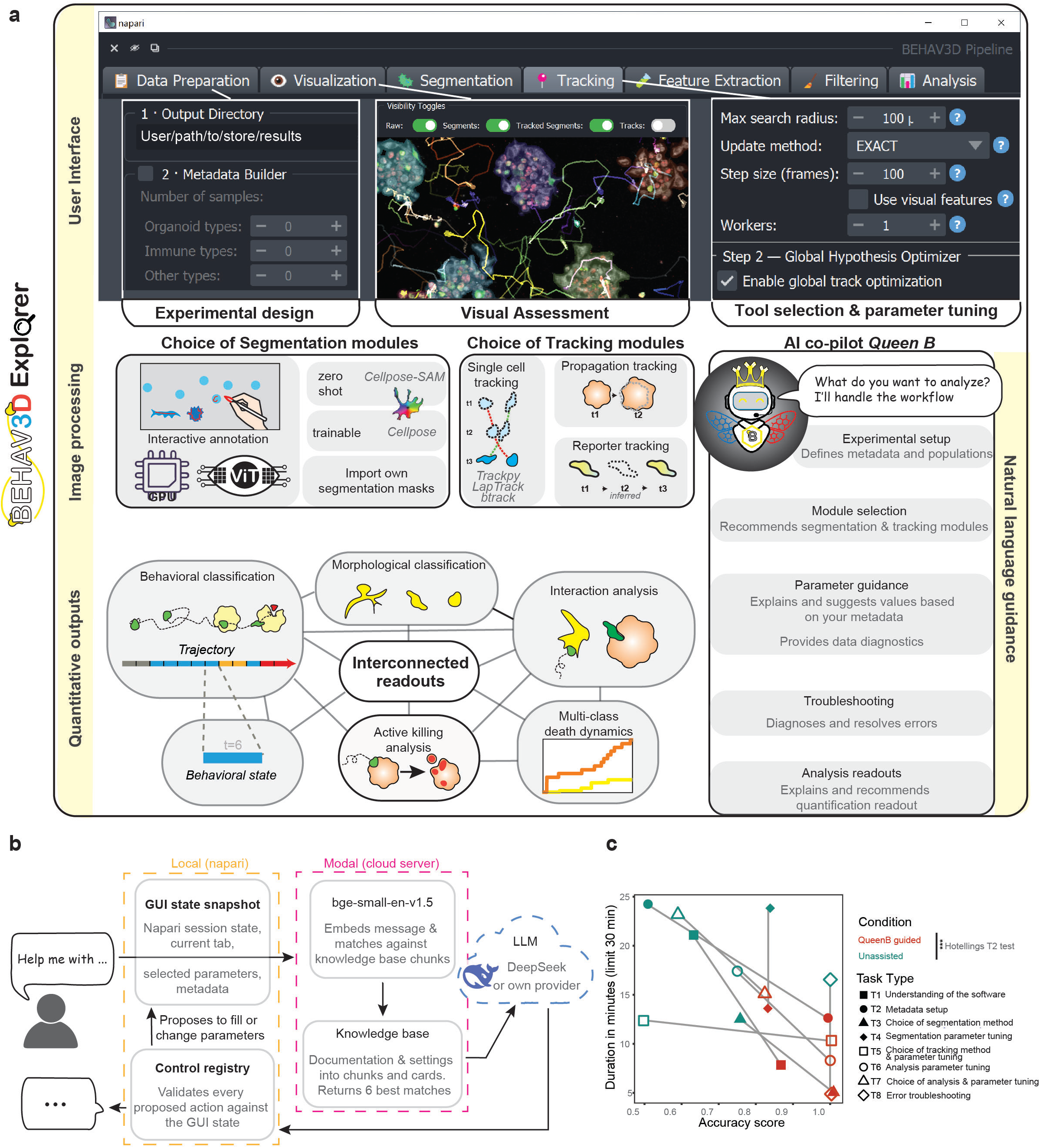
Workflow of BEHAV3D-Explorer and architecture and performance of QueenB assistance. **(a)** Overview of the BEHAV3D-Explorer graphical user interface, image processing modules and main analysis outputs. **(b)** Architecture of the QueenB natural-language assistant. **(c)** Task-completion time and accuracy for BEHAV3D-Explorer tasks performed with (red) and without QueenB assistance (green). One-sample Hotelling’s *T*² test on per-task differences, *p* = 3.63 × 10⁻⁴, n=8 tasks.

BEHAV3D-Explorer is implemented as a standalone napari-based application with a no-code installer, supporting common microscopy formats (**Fig. 1a**). Its modular architecture accommodates diverse experimental designs through flexible metadata definition and interchangeable segmentation and tracking methods. Segmentation options include the foundation model CellposeSAM^11^ and vision transformer-based segmentation methods (ConvPaint^12^), alongside alternative approaches optimized for specific trade-offs in accuracy, speed and channel bleed-through outweighed against computational costs (**Extended Data Fig. 1**, **Supplementary Note 1**). Tracking strategies extend beyond conventional motile-cell tracking to accommodate tissue fragmentation during cell death (**Extended Data Fig. 2a**), densely interacting cells (**Extended Data Fig. 2b**) and temporally fluctuating reporter activity (**Extended Data Fig. 2c**). Together, these modules allow analysis workflows to be tailored to the technical and biological heterogeneity of live-imaging datasets. Users can inspect and correct segmentation masks and cell tracks, and visually assess analytical outputs directly against the original imaging data (**Fig. 1a**).

Because deciding which combination of modules and parameters suits a given dataset is often the main hurdle for non-expert users, we integrated QueenB, a natural-language assistant that guides users through the analysis process (**Fig. 1b**, **Supplementary Video 1**). By describing their dataset, analysis challenge or biological objective, users receive context-specific recommendations for metadata configuration, segmentation and tracking strategies, parameter optimization, downstream analyses and troubleshooting. QueenB can also generate editable analysis configurations, which users can inspect and modify before execution (**Extended Data Fig. 3**). Across eight representative workflow tasks, QueenB improved beginner-user performance, enabling faster task completion (2.13-fold on average) and/or greater agreement with expert-defined workflows (94,3% with QueenB assistance compared to 69% without assistance; **Fig. 1c**, **Supplementary Note 2**).

Importantly, beyond lowering the implementation barrier, BEHAV3D-Explorer integrates multiple informative readouts that are usually quantified separately, which we demonstrate across experimental conditions, spanning *in vitro* organoid co-cultures, intravital imaging and *ex vivo* tissue explants (**Extended Data Fig. 4a**, **Supplementary Table 2**). Patient-derived organoids (PDOs) are used to evaluate immune-cell therapies within increasingly complex 3D tumor contexts ^2,3,13–16^, yet extracting meaningful readouts from these immune-organoid co-cultures requires resolving several cell populations and their behavior over time and how it relates to tumour targeting. Using the metadata framework, BEHAV3D-Explorer assigns multiple target populations, in this case healthy breast organoids and breast cancer PDOs, (**Fig. 2a**, **Supplementary Video 2**) and differentially quantifies their response to the same treatment. This demonstrates preferential targeting of tumor organoids, whereas healthy organoids were largely spared (**Fig. 2b**), offering a combined readout of on-and off-tumor activity, essential for toxicity profiling.

**Figure 2.**
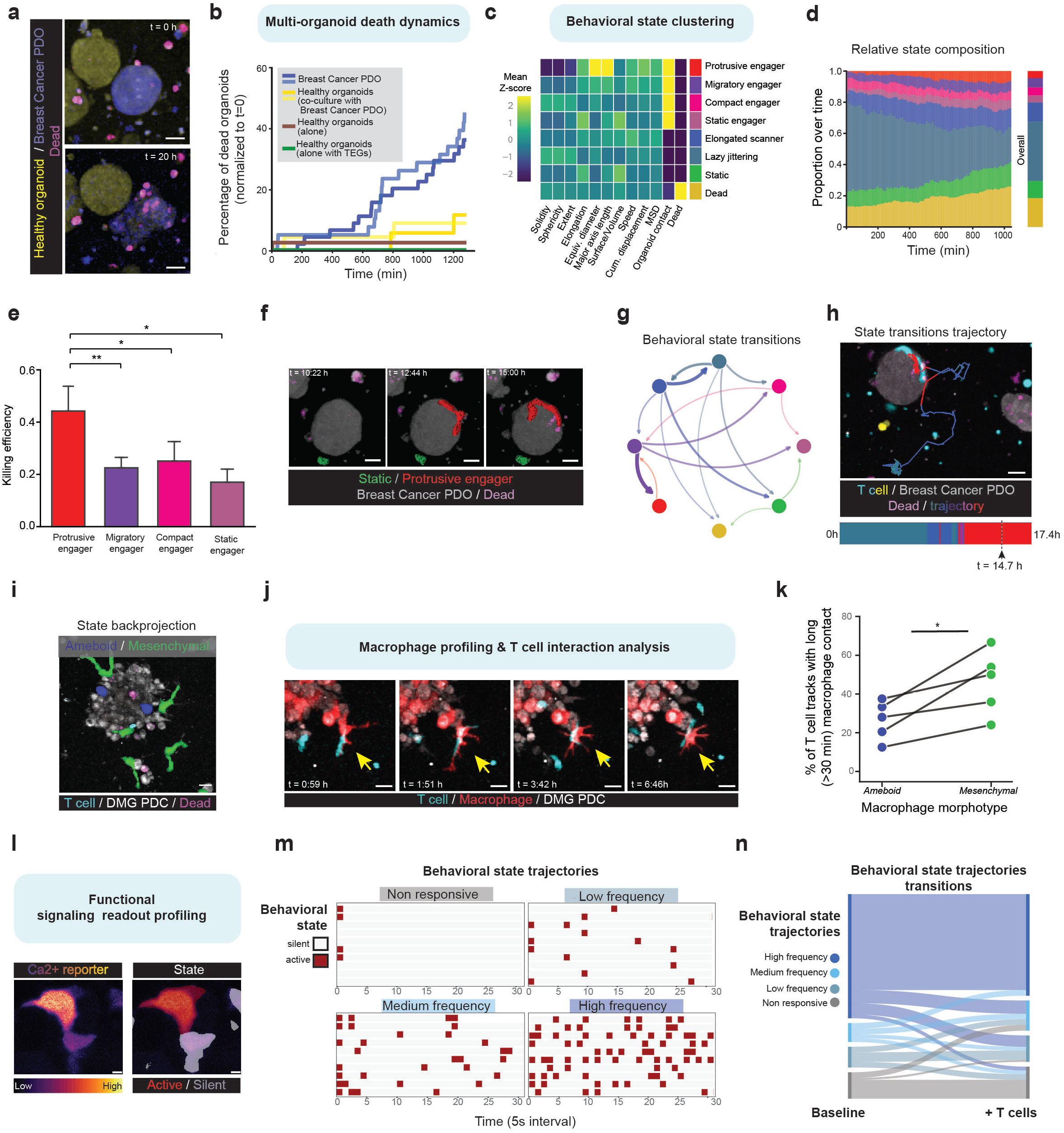
BEHAV3D-Explorer enables complex biological readouts across diverse live-imaging datasets. **(a,b)** Representative example (a) of co-cultures of healthy and patient-derived breast cancer organoids exposed to engineered T cells and (b) quantified survival dynamics **(c)** Mean z-score of state-defining features per behavioral state. **(d)** Relative abundance of each behavioral state over time (left) and overall state distributions across all time points (right). **(e)** Bar plots showing per-cell killing efficiency; linear mixed-effects model with imaging well as a random effect (5 imaging wells). P versus protrusive engager: migratory=0.002, compact=0.021 static=0.038; all other comparisons not significant. **(f)** Time-lapse frames of a representative protrusive engager T cell. **(g)** Transitions between behavioral states (minimum state duration 3 timepoints, transitions >20% shown). **(h)** State sequence of a single T cell backprojected over the full timelapse. **(i)** Representative example of morphological profiling identified distinct macrophage phenotypes in co-cultures with DMG PDCs and CAR T cells. **(j)** Representative time-lapse frames of multi-hour T cell–macrophage contact. **(k)** Proportions of T cells entering in long contact (>30 minutes) with mesenchymal versus amoeboid macrophages (two-sided paired t-test, P=0.025, n=5 imaging wells). **(l)** Representative calcium reporter images of pancreatic islet cells (left) and backprojected states (right). **(m,n)** Representative behavioral state trajectories (m) and their transitions after allogeneic CD8+ T cell addition (n). All scale bars, 20 μm.

We next investigated the underlying T cell behavior associated with tumor targeting through single-cell tracking and unbiased behavioral classification, using multicolor labeling to facilitate accurate tracking (**Supplementary Video 3**). Trajectory-level analysis recovered previously described T cell subsets^2,3^, including ‘super-engagers’ characterized by prolonged organoid contact, potent serial-killing capacity and substantial morphological plasticity, previously observed only qualitatively (**Extended Data Fig. 4b,c**). BEHAV3D-Explorer now enables quantitative profiling of T cell behavior at individual timepoints, identifying eight distinct morphodynamic states whose relative abundance and temporal dynamics can be quantified (**Fig. 2c-d**). By coupling these states to the newly implemented active-killing analysis, which links T cell-organoid contacts to subsequent increases in organoid death signal, we identified the ‘protrusive engager’ state as most strongly associated with tumor killing, enabling quantitative characterization of the dynamic behaviors and cytotoxic activity underlying the previously described ‘super-engager’ phenotype^3^ (**Fig. 2e,f**). Furthermore, state-transition analysis quantified how T cells switch between behavioral states, enabling reconstruction of the behavioral routes leading to this killing-associated state. This revealed even rare, complex behavioral sequences progressing from ‘lazy jittering’ cells through ‘elongated scanners’ and ‘migratory engagers’ to ‘protrusive engagers’. Importantly, these rare behavioral routes could not only be quantified but also directly mapped back onto the original imaging data (**Fig. 2g,h**)), revealing transitions previously obscured by trajectory-level classification.

To demonstrate additional analytical capabilities in a more complex multicellular microenvironment, we applied BEHAV3D-Explorer to co-cultures of macrophages, chimeric antigen receptor (CAR) T cells and diffuse midline glioma patient-derived cells (PDCs) (**Supplementary Video 4**), linking macrophage morphological heterogeneity to distinct patterns of cell-cell interactions. Myeloid cells, such as macrophages, regulate CAR T cell activity and can contribute to immunosuppression^17,18^, yet how their dynamic morphology shapes these interactions remains poorly characterized. Morphodynamic profiling revealed two previously described macrophage morphotypes^19^, amoeboid and mesenchymal-like (**Fig. 2i**), which differed in their propensity to establish prolonged contacts with CAR T cells (**Fig. 2j,k**), an interaction pattern implicated in regulating T cell function^20^. Beyond *in vitro* systems, we reanalyzed published intravital recordings of the popliteal lymph node during HIV infection^21,22^ (**Supplementary Video 5**). Jointly profiling independently tracked uninfected and HIV-infected CD4+ T cells revealed fewer fast-migrating behaviors among infected cells (**Extended Data Fig. 4 d**), consistent with the reduced motility reported in the original study^21^. Lastly, we extended the analysis to functional reporter imaging of a mouse pancreatic islet explant expressing the Ca²⁺ reporter GCaMP3, before and after addition of allogeneic CD8+ T cells, modeling immune-mediated graft injury (**Fig. 2l**, **Supplementary Video 6**). Using our custom reporter-propagation tracking (**Extended Data Fig. 2c**), state and trajectory profiling resolved heterogeneous Ca²⁺ activity at baseline (**Fig. 2m**) and its changes after T cell addition (**Fig. 2n**), as a measure of islet function. Together, this demonstrates how BEHAV3D-Explorer connects high-level multimodal imaging readouts within a single user-accessible analytical framework across diverse experimental settings.

## Supplementary information

**Extended Data Figure 1. Segmentation Benchmarking of BEHAV3D Explorer**. **(a)** Wall-clock time (s) versus peak RAM (GB) during training of each segmentation option for T cell (top) and organoid (bottom) segmentation. (b) Wall-clock time (s) versus peak RAM (GB) during inference of each segmentation option for T cell (top) and organoid (bottom) segmentation. Marker shape, size and fill encode method and annotation budget. Large markers represent the mean of n=3 repeats; small translucent markers represent individual runs. Time is log-scaled, peak RAM (GB) is linear.

**Extended Data Figure 2. Custom-designed tracking methods for organoid fragmentation, near-static/plastic cell tracking, and reporter signaling monitoring**. **(a)** Schematic representation of fragmentation-propagation tracking, which preserves object identity during organoid deformation, death, and fragmentation. **(b)** Schematic representation of bounded-propagation tracking for dense, slowly moving, and plastic cell populations. Cell identities are propagated across consecutive timepoints based on spatial overlap, preserving tracks despite substantial shape changes and transient cell-cell contacts. **(c)** Representative GCaMP3 calcium-reporter imaging time course (top), corresponding segmentation results showing intermittent detection failures caused by signal fluctuations (middle), and reporter-propagation tracking masks (bottom). This approach maintains object identity and enables longitudinal quantification of reporter activity despite transient signal loss. Scale bar, 30 μm.

**Extended Data Figure 3. Example chats demonstrating QueenB capabilities**. Representative examples of QueenB-assisted interactions during image-analysis workflow design, including metadata configuration, module selection, parameter recommendation, troubleshooting, workflow adaptation to new datasets, module explanation, analysis recommendation, and interpretation of analytical outputs. QueenB provides context-aware guidance by leveraging dataset metadata, analysis results, and built-in knowledge of BEHAV3D-Explorer workflows.

**Extended Data Figure 4. Experimental datasets analyzed with BEHAV3D-Explorer, including trajectory level and state behavioral profiling. (a)** Schematic overview of the 5 experimental use-cases of BEHAV3D-Explorer. **(b)** UMAP representation of 9 color-coded behavioral clusters from trajectory-level profiling of 1,190 T cell tracks. **(c)** Backprojection of selected behavioral subtypes onto a representative field of view. **(d)** Behavioral state profiling reveals motility changes in HIV-infected versus uninfected T cells (two-sided paired t-test, n=3 fields of view; bars show mean ±95% CI; P=0.024, 0.081 and 0.097 for fast, slow and static states respectively).

**Supplementary Note 1.** Explanation of segmentation backends and their benchmarking.

**Supplementary Note 2.** QueenB evaluation study design.

**Supplementary Table 1.** Feature comparison of BEHAV3D-Explorer against contemporary live-imaging analysis software.

**Supplementary Table 2.** Details of the five case-study datasets used to showcase BEHAV3D-Explorer analysis capabilities.

**Supplementary Video 1.** BEHAV3D Explorer workflow overview including QueenB assistance.

**Supplementary Video 2.** Representative 3D time-lapse of the dual healthy and breast cancer organoid co-culture experiment.

**Supplementary Video 3.** Representative 3D time-lapse of the multicolor T cell co-culture experiment with T cell segmentation and tracking.

**Supplementary Video 4.** Representative 3D time-lapse of the macrophage and T cell with DMG PDC co-culture experiment.

**Supplementary Video 5.** Representative 3D time-lapse of the intravital microscopy experiment (uninfected and HIV infected T cells in popliteal lymph node).

**Supplementary Video 6.** Representative 3D time-lapse of the pancreatic islet Ca²⁺ imaging experiment.

## Methods

### BEHAV3D-Explorer implementation

BEHAV3D-Explorer is implemented in Python (3.12) and distributed as a *napari* plugin, so that three-dimensional rendering and every analytical step share a single process and canvas. The interface is built with Qt (via *qtpy* and *magicgui*) and organized into seven tabs that follow the analysis workflow in order (Data Preparation, Visualization, Segmentation, Tracking, Feature Extraction, Filtering and Analysis) together with a processing queue and a co-pilot assistant panel that span all tabs. The Data Preparation tab holds the single source of truth for a project: the experiment metadata table, the output directory and a parameter dictionary, serialized to YAML after every change, which all other tab references. Long-running computation runs on background threads with progress reported to both the plugin and the native *napari* activity dock. Every processing step saves its output using the same consistent file and folder naming pattern. The platform can be installed either as a Python package, or via standalone installers for Windows, macOS and Linux that require no coding experience. These installers automatically set up the full working environment and detect whether the computer has a CUDA-compatible GPU, Apple Silicon, or only a CPU, installing the correct version of *PyTorch* accordingly.

### Data management

BEHAV3D-Explorer reads microscopy datasets stored in CZI, LIF, TIFF, and Zarr/OME-Zarr formats through a common loader and converts them to chunked Zarr arrays, with one chunk per timepoint, giving lazy, out-of-memory access to volumetric time-lapse datasets via Dask. Axis order is read from file metadata where it is encoded or is otherwise requested from the user and then normalized to a canonical TCZYX layout. An experiment is defined in an editable metadata table, one row per field of view, built through a graphical metadata builder. Users declare any number of biological entities (organoid-like, immune-like and other cell populations) and name them freely. Sample-level fields record voxel spacing in xy and z, frame interval, experimental conditions, the raw image path and an optional death-reporter channel. Optionally, immune populations imaged across multiple channels are expanded into per-channel types for acquisition and recombined into a single merged population for analysis.

### Segmentation modules

BEHAV3D-Explorer integrates five segmentation backends behind one interface: a zero-shot deep-learning model (Cellpose-SAM), a trainable deep-learning workflow (CellPose), and three complementary pixel-classification workflows trained from sparse annotations (a CPU-only classifier, a GPU-accelerated random-forest classifier (APOC) and ConvPaint). Externally generated segmentations can also be imported.

Cellpose-SAM^11^ is used as a zero-shot backend requiring no manual annotation. Because it cannot share dependencies with the Cellpose v.3 backbone used elsewhere in the platform, it runs out of process in an isolated environment, with progress streamed through a JSON-Lines interface so reporting matches the in-process backends.

Cellpose v3^23^ is available for datasets requiring improved boundary accuracy or separation of touching objects and is run natively in three dimensions using each sample’s own anisotropy, with minimum object sizes set by cell-type category. Users can apply existing pretrained models directly or fine-tune them on manually annotated data using an accompanying notebook. Death masks are generated separately by Otsu thresholding of the death-reporter channel.

For the user-trainable pixel classifiers, APOC^24^ is the default pixel-classification workflow, chosen for its broad hardware compatibility and speed. Sparse foreground and background annotations, drawn by the user in napari on timepoints sampled across all samples of an experiment, train a GPU-accelerated random-forest classifier from multiscale intensity, edge and texture features; a CPU-only implementation is also provided that computes the same features at physically calibrated scales. For both, a model is trained for each cell type separately. Annotations and training data are stored alongside the classifier, allowing incremental refinement across experiments.

For datasets requiring broader visual and contextual information, ConvPaint^12^ combines a choice of pretrained feature extractors (for example VGG-16, DINOv2, or Cellpose/Ilastik backbones) with a gradient-boosted classifier; one unified multiclass model spans all declared cell types and is evaluated plane by plane through z to bound memory.

Because both pixel-classification routes return per-class probabilities, instances are recovered by a shared, anisotropy-aware seeded watershed on the distance transform (or on the class-probability map), followed by hole filling, morphological opening and size filtering; flat, single-plane artefacts are also removed. Populations imaged across multiple fluorophores are reconciled by resolving shared voxels to the largest overlapping object.

### Tracking modules

BEHAV3D-Explorer integrates six tracking backends with one output format: a track table and a tracked label image in which each object carries its track identity throughout the movie, allowing tracks to be overlaid on the raw data and validated before analysis. For freely migrating cells, three backends adapt established particle-tracking methods: nearest-neighbor linking with an adaptive search radius (TrackPy), linear-assignment-problem tracking with gap closing (LapTrack), and Bayesian tracking with a Kalman motion model (btrack). Because these tracking approaches do not perform reliably across all biological scenarios, especially when identity across time if better defined by spatial overlap than by centroid displacement, we developed three custom tracking backends: (1) fragmentation-propagation tracking, which preserves individual organoid identities through the deformation and fragmentation that accompany cell death (**Extended Data Fig. 2a**); (2) bounded-propagation tracking, designed for dense, slowly moving, and interacting cell populations (**Extended Data Fig. 2b**); and (3) reporter-propagation tracking (**Extended Data Fig. 2c**), tailored for intermittent signals, such as fluorescent reporters, that cannot be segmented reliably at every timepoint.

#### Fragmentation-propagation tracking

This method relies on objects being correctly separated and non-fragmented at t=0, as here the total number of objects is set. This allows the tracking to correctly separate touching objects that lost clear boundaries or assign fragments to organoid identities (**Extended Data Fig. 2a**). At t=0, tracks are initialized directly from the segmentation labels of that timepoint. For each consecutive timepoint pair (t, t+1), after, the track labels at t are overlaid onto the foreground mask at t+1 and labelled pixels falling outside this mask are removed. A marker-controlled watershed, seeded from the remaining track labels retained from t, is applied to relabel t+1. To connect fragmented segments, a second watershed pass is applied, where the foreground mask is now dilated by a fixed margin (2 voxels) to connect fragmented foreground mask. The output is then clipped back to the original t+1 foreground mask. Objects below a minimum size (default 100 voxels) are removed after each timepoint. Any objects newly appearing after t=0 without a track label after these watershed steps are not separately instantiated.

#### Bounded-propagation tracking

This method enforces that a single-track identifier can never span more than one spatially disconnected object. Like fragmentation-propagation tracking, it allows for touching objects to be correctly separated, while enabling for the creation of new tracks outside of t=0 (**Extended Data Fig. 2b**). At t=0, tracks are initialized directly from the segmentation labels of that timepoint. For each consecutive timepoint pair (t, t+1), the track labels at t are overlaid onto the foreground mask at t+1 and labelled pixels falling outside this mask are removed. Before watershed, the foreground mask at t+1 was independently partitioned into connected component regions using full voxel connectivity. Each track at t was then assigned to the region its previous footprint overlapped most by pixel count and overlapping pixels with other regions or background are removed. Regions claimed by no track are left unflooded or are assigned a newly instantiated track. Resulting tracked objects are filtered by a user-defined size filter (default 20) and a 2D object filter to remove any untrue 3D objects that only occur in a single plane.

#### Reporter propagation tracking

This method works on near-static objects whose segmentation is intermittently absent because of a fluctuating reporter signal (e.g. a calcium reporter), when no stable cell label is available. Instead of frame-to-frame linking, all segmented instances detected across time are first pooled and objects below a minimum size (default 100 voxels) are discarded as noise. The remaining instances are then grouped based on spatial overlap, by at least a minimum fraction (default 10%), with the largest instance currently assigned to each group, and all comparisons made against this single representative reference mask. For each group, the largest instance observed throughout the time lapse is selected as the canonical mask and propagated unchanged to all timepoints, generating a continuous full-duration track for every object (**Extended Data Fig. 2c**).

### Multicolor segmentation and tracking

To make dense populations easier to segment and track, cells can be labelled with several fluorophores simultaneously, so that each fluorophore and their unique combinations divide the total population into less dense subpopulations. In BEHAV3D-Explorer, a cell type can be defined as multicolor, and the following workflow then adapts to this set-up. Each subpopulation is first segmented separately, and then overlapping segments between them are resolved: when voxels are claimed by more than one segment, they are assigned to the largest overlapping segment and the overlap is removed from the smaller ones. The remaining part of each smaller segment is eroded by one voxel (by default). Segments removed by this erosion are discarded as residual fragments, while the rest keep their original non-eroded shape. Each resolved subpopulation is then tracked separately, so links are only made between cells of the same subpopulation. Finally, the tracked segments are merged into a single population, with each track’s source subpopulation kept as an annotation. All further analysis steps are run on this merged population.

### Feature extraction

A configurable set of morphological, intensity, motility, contact, invasiveness and viability descriptors can be computed for every tracked object at every timepoint, in physical units, for all declared populations, and users select which descriptors to extract for a given analysis. Morphology is calculated using *regionprops* (scikit-image, voxel-spacing aware), with shape/orientation metrics derived from per-object voxel-based calculations and principal axes. Two thresholds are set by the user: a contact threshold, which next to direct physical contact defines a distance within which neighboring objects are additionally recorded as being in contact (evaluated both as pixel adjacency and against the physical threshold), and a death threshold, above which an object is called dead (see Organoid death dynamics analysis, below). Death is measured as the fraction of a segment’s voxels overlapping the segmented death-reporter mask. For organoid types, dead voxels inside other cell types are masked out. Missing timepoints within a track are interpolated and flagged, and tracks are filtered on length, size and/or viability at the first timepoint before analysis.

### Quantitative analysis

#### Organoid death dynamics analysis

Organoid death is quantified as the fraction of an organoid’s segmented voxels overlapping a dead-cell dye mask, excluding dead-mask voxels that coincide with other segmented cells. An organoid is classified as dead once this fraction exceeds a user-defined threshold, and the call is retained at all later timepoints regardless of fluctuation. Each track is scored per timepoint as alive, dead or disappeared, with the last flagging tracks that end before the observation window closes without ever crossing the death threshold (likely fusion or segmentation loss). State counts are aggregated per sample and timepoint, normalized to the starting organoid count, and reported as mean ± s.e.m. across tracks, per sample or pooled by condition.

#### Interaction analysis

Contact between cells is defined from the segmentation-based proximity computed during feature extraction. For each organoid, contact with a given effector population accumulates as a running count of timepoints in contact, and each organoid’s terminal fate allows contact histories to be compared between organoids that die and those that survive, per sample or pooled by condition. Because organoids die at different times, contact is also examined relative to time of death: contacts within a user-defined window before death (default 60 min) are attributed to that event and averaged across tracks as mean ± s.e.m.

#### Active killing analysis

Active killing attributes a target population’s overall rise in death signal to individual effector cells by flagging the timepoints at which an effector touches a target that shows a death signal increase large enough to count as a killing event.

The detector first identifies contact events like in the *Interaction analysis*. An event has to last at least the minimum contact duration to be considered (default 1 timepoint). Then, at every timepoint of that contact *t*, it evaluates each touched target independently: it reads that target’s own death signal at the current timepoint (*D_start_*) and again at a tunable ‘Observation window’ *W* (default 5 timepoints) later (*D_end_*), and takes the difference (*ΔD*):

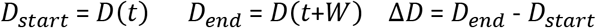

This increase ΔD is compared to a threshold θ, which is either an absolute increment in the death signal (*θ_abs_*) or a fold-change *m* (default 1.5) over the target’s own signal at contact timepoint *t* (*θ_fc_*). An effector is flagged as actively killing when its targets death signal rise exceeds the threshold, and the killing efficiency is the ratio of the observed rise *ΔD* to the threshold *θ*:

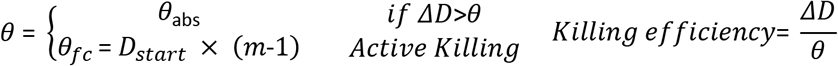

Each target is judged only against its own signal, with no cross-target or sample-wide averaging. When an effector touches several targets at once, the one that clears the threshold by the largest margin is reported as that timepoint’s target. The observation window and threshold are set based on the imaging interval and the expected biology rather than fixed values.

#### Behavioral state classification

To capture how a cell is behaving at each moment, per-timepoint behavioral states for each cell at every timepoint are inferred in a multi-step pipeline. First, a Gaussian hidden Markov model (*hmmlearn*) is fit based on a set of user-defined features. These include per-timepoint features (optionally smoothed and log1p-scaled) and features computed over a user-defined trailing window (default N timepoints; e.g. net displacement), which set the timescale of the captured dynamics. All features are standardized using z-score normalization. The first timepoint of each track is excluded from fitting, since dynamic features default to zero here, and later backfilled to allow every timepoint to have an HMM state. The model is then fit on all tracks with a user-defined fixed number of states. Binary or categorical features (e.g., death status, contact, active killing) are usually excluded from model fitting and instead used post hoc to subdivide each inferred HMM state into behavioral clusters. These clusters can be manually renamed or merged after visual inspection to their final behavioral state labels. Per-cell, per-timepoint state assignments were then used to generate state-composition and state-transition summaries across samples or condition groups. Furthermore, the full processing pipeline (including feature selection, preprocessing parameters, fitted HMM and cluster renaming/combination schedules) are saved in a single .pkl file, allowing reuse on new data as a classification pipeline.

#### Behavioral trajectory classification

While behavioral states describe individual timepoints, behavioral trajectory classification groups the behavior of cell tracks. In BEHAV3D-Explorer this can be done in two ways, feature-based or state-based. For feature-based trajectory clustering, each track is represented as sequences of user-selected single-timepoint features over the track and dynamic time warping is applied to calculate distances between all tracks. For state-based trajectory clustering, tracks are clustered by the overall pattern of states they pass through across the entire movie. Each track is represented as its sequence of behavioral-state labels, and pairwise distances can be computed in two ways, using multivariate dynamic time warping (dtaidistance) or trajectory-level summary features (per-track state proportions, per-state bout statistics and state transitions) to quantify how similar two cells’ sequences of behavior are. For both feature-based or state-based distance calculation, tracks can then be grouped by either agglomerative hierarchical or Leiden clustering, yielding trajectory classes that represent distinct trajectory behavioral phenotypes. Like behavioral states, the user can manually rename/combine the resulting clusters after visual inspection for resolving final track classifications. Trajectory assignments can then be used to produce composition and diagnostic reports. A random forest classifier (scikit-learn), trained on trajectory-level summary features allows these labels to be reapplied to new tracks or datasets.

### QueenB natural-language assistant

QueenB is a natural-language assistant integrated into the BEHAV3D-Explorer interface. Users describe a biological question or analysis goal in plain language, and QueenB responds with a recommended workflow, parameter suggestions, or an editable analysis configuration. Its responses come from two sources: a knowledge base built from the platform’s documentation and parameter definitions, and the live state of the user’s current session, including loaded metadata, workflow progress and available interface controls, so that suggestions are grounded in what is actually loaded and possible to do at that point.

QueenB’s knowledge base is built from the platform’s documentation, the explanatory notes in its code, and a catalogue of every configurable parameter (its workflow step, cell category, data type, default value, allowed values and explanation), generated automatically from the platform’s own settings and feature definitions. Documentation is split into heading-anchored chunks, and each parameter card forms its own chunk; all chunks are embedded with a 384-dimensional sentence-transformer (BAAI/bge-small-en-v1.5) and indexed for cosine-similarity retrieval. For each query, the embedded user message and current workflow step retrieve the most relevant chunks, which are combined with deterministic, step-specific guidance cards.

QueenB runs as a separate panel within the napari interface, connected to BEHAV3D-Explorer through a live link that keeps both sides in sync. Before each exchange, QueenB sends a snapshot of the full session: the active workflow tab, selected method and cell type, loaded or in-progress metadata, etc. Requests run in the background so the interface stays responsive, and the assistant’s response is displayed as it is generated rather than all at once. The conversation and the live session snapshot are sent to a backend service (FastAPI on a CPU-only Modal deployment). Deterministic rules answer routine requests directly. Otherwise, the service retrieves relevant knowledge, builds an instruction set that treats the live interface state as ground truth and limits the assistant to the actions and controls currently available, and queries a large language model, configured by default to favor consistent, predictable responses over creative ones, to generate a reply that is streamed back to the interface as it is produced. QueenB runs by default on DeepSeek (deepseek-v4-flash) through the BEHAV3D assistant service. Alternatively, users can select a different model provider (OpenAI, Anthropic, Google Gemini, Mistral or DeepSeek) with their own API key, which is stored encrypted in their operating system’s keychain and sent only to that provider, never to the BEHAV3D service; document retrieval and checks on proposed actions remain on the service. Each proposed action, such as filling metadata fields, updating a control, navigating tabs, queuing a processing step, opening a preview, or creating a cell-type group, is validated against the live control registry for identity, data type, allowed range and current enabled or visible state, and no-op requests are discarded. Read-only actions (previews, navigation) apply immediately; actions that change a value, queue new work or create a file require explicit user confirmation before being applied. A deterministic local wizard handles routine sample and cell-type metadata setup without involving the language model, and if the hosted service is unavailable, QueenB falls back to answering parameter questions from its local parameter definitions only.

### Benchmarking of QueenB

QueenB performance was evaluated in a within-subject, counterbalanced usability study with github commit 9590261. Eight participants with a biology background, but no prior experience with BEHAV3D-Explorer or advanced image analysis, each completed the same eight workflow-configuration tasks on a four-channel organoid-T-cell co-culture dataset, covering software understanding, metadata definition, segmentation-method selection, segmentation-parameter tuning, tracking configuration, active-killing analysis setup, behavioral-state analysis setup and error diagnosis (Tasks described in **Supplementary Note 2**). Four tasks per participant were completed with QueenB available (assisted) and four using the online Github documentation and wiki only (unassisted), assigned so that each task had the same number of assisted and unassisted tries, giving 64 task-instances. Maximum tasks time was capped at 30 min. Each submitted task configuration was scored by three independent expert moderators as full, partial or no agreement. Configuration time, agreement with the gold standard, and self-reported confidence (1-10) were recorded for each task. Task instances affected by factors unrelated to the experimental condition were excluded due to temporary chatbot unavailability caused by a transient DeepSeek backend outage, misinterpretation of the task objective, incomplete task briefing by the moderator, or a task not completed (participant had to leave the session). To maintain a balanced study design, the assignment of conditions for subsequent participants was adjusted when exclusions occurred, ensuring a final dataset containing at least three assisted and three unassisted observations per task. Per-participant results are provided in **Supplementary Note 2**.

### Imaging datasets

BEHAV3D-Explorer was applied to a variety of live imaging datasets newly generated or from previously published repositories summarized in **Supplementary Table 2**. Newly generated datasets are described below.

#### Multi-organoid breast cancer dataset (Dataset 1)

Healthy breast organoids and patient-derived breast tumor organoids were co-cultured with T cells engineered to express a gamma delta T cell receptor (TEGs) within the same imaging well or in separate wells. Organoids and TEGs were prepared as previously described^2,3^. For these experiments, 40.000 T cells per condition were stained using Celltracker green CMFDA (Thermo Fisher) 1:5000 in PBS for 10 minutes at 37 °C. Healthy organoids were stained using Celltracker orange CMRA (Thermo Fisher) 1:4000 and cancer organoids in Celltrace violet (BD Horizon) 1:4000 both in PBS for 15 minutes at 37 °C. For each imaging well, 1:800 TO-PRO3 dead dye (Thermo Fisher Scientific) was added to capture cell death. Live cell imaging was performed on the LSM Zeiss 980 confocal laser scanning microscopes (Carl Zeiss Microscopy, Jena, Germany) equipped with an environmental incubation chamber at 37 °C and 5% CO2. Images were acquired using a 20x Air objective with a zoom of 0.6. Optical Z-stacks were captured over a total thickness of 100 µm and a time interval of 2 minutes at a resolution of 1.77 μm in X Y Z.

#### Multicolor T cell and breast cancer organoids dataset (Dataset 2)

Co-cultures of B7-H3 CAR CD8+ T cells and patient-derived breast cancer organoids were prepared as previously described^2,3^. Breast cancer PDOs were stained with CellTracker Green CMFDA (1:2500, Thermo Fisher Scientific) in +++ media for 20 min at 37°C. T cells were stained in PBS for 20 min at 37°C using three combinations: Only eFluor™ 450 (1:1000, Thermo Fisher Scientific), only CellTracker Orange (1:2000, Thermo Fisher Scientific) or both, providing three differently stained T cell fractions. In a 96-well glass-bottomed imaging plate (Cellvis), 8.000 T cells (equal proportions of each of the three differently stained T cells) were combined with each PDO line per well in 50% breast cancer PDO, 50% T cell media (RPMI gultamax, 1% penstrep, 10% FCS) with 2.5% BME and 1:1000 TO-PRO-3 dead dye (Thermo Fisher Scientific). Live cell imaging was performed on the Zeiss 980 confocal laser scanning microscope (Carl Zeiss Microscopy, Jena, Germany) equipped with an environmental incubation chamber at 37°C and 5% CO2. Images were acquired with a 20x air objective with 0.6 zoom, Optical z-stacks were captured over a total thickness of 50 µm with a time interval of 2 min and 1.38 µm voxel size.

#### Diffuse midline glioma (DMG)-macrophages-CAR-T-cell co-culture dataset (Dataset 3)

Three DMG PDC lines were cultured as previously described^25–28^ and used for co-culture experiments 2-4 days after passaging. CD14+ monocytes from healthy donors were differentiated into naïve macrophages using 10ng/ml of GM-CSF for 24 hours followed by overnight equilibration in medium without GMCSF. GD2 CAR T cells were generated and expanded as previously described^29^. Macrophages and DMG PDCs were co-cultured for 72 h before addition of GD2 CAR T cells. For live-cell imaging, macrophages were labelled with CellTracker Green, DMG PDCs with CellTracker Red, GD2 CAR T cells with eFluor 450, and dead cells with Annexin VI-CF555. Imaging was performed on a Zeiss LSM880 confocal microscope (Carl Zeiss Microscopy, Jena, Germany) equipped with a 37°C, 5% CO₂ incubation chamber using a 20×/0.8 NA Plan-Apochromat objective. Time-lapse Z-stack images (50 μm total depth) were acquired every 131 seconds at a resolution of 1.77 μm in X Y Z.

#### Calcium imaging of pancreatic islet tissue explants dataset (Dataset 5)

WT and DBA/2 mice were purchased from Charles River Laboratories. Experiments were conducted using age-matched mice of both sexes aged 15-22 weeks. Animals were housed in a temperature-controlled room (22 °C, 50–60% humidity) under a standard 12-hour light/dark cycle. All experimental procedures strictly complied with local state regulations (Landesamt für Verbraucherschutz, Saarland; approval numbers: AZ 2.4.1.1 and 11/2021). Splenocytes were harvested from 15-to 22-week-old donor mice as described previously^30^. Naive CD8+ T cells were positively isolated from WT mice using the Dynabeads FlowComp Mouse CD8 Kit (Invitrogen) according to the manufacturer’s instructions. Isolated CD8+ T cells were stimulated with anti-CD3/anti-CD28 activation dynabeads at a 1:0.6 cell-to-bead ratio and cultured at a density of 1 x 106 cells/mL in 24-well plates. Cells were maintained in RPMI medium (Invitrogen) supplemented with 10% FCS, 1% penicillin/streptomycin (Invitrogen), and 50 µM 2-mercaptoethanol. On day 2 post-stimulation, 100 U/mL recombinant mouse IL-2 (Gibco) was added to promote T-cell expansion. The resulting activated effector T cells were harvested on day 4 for co-culture experiments. Primary pancreatic islets were isolated as previously described^31,32^ from DBA/2 mice. Isolated islets were seeded into 8-well glass-bottom imaging chambers (Sarstedt) and maintained in RPMI 1640 medium containing 10% FCS and 10 mM HEPES at 37 °C in a humidified 5% CO2 incubator. To express the cytosolic calcium indicator GCaMP3, islets were transduced with GCaMP3-encoding lentivirus 1 day post-isolation. Transduced islets were cultured for 8-10 days post-isolation to ensure stable adherence to the glass surface and adequate GCaMP3 expression prior to live-imaging assays. Live-cell imaging was performed on a Zeiss LSM 780 confocal laser scanning microscope (Carl Zeiss Microscopy, Jena, Germany) equipped with an environmental incubation chamber maintained at 37 °C and 5% CO2. Images were acquired using a 40x Plan-Apochromat oil-immersion objective (NA 1.3). Optical Z-stacks were captured over a total thickness of 10 μm at 2 μm step intervals (6 slices per stack) with a frame size of 1024 x 1024 pixels and a spatial resolution of 0.33 μm x 0.33 μm per pixel. GCaMP3 fluorescence dynamics in individual islet cells and brightfield were recorded with a 5 seconds time interval during 32 timepoints before and after effector CD8+ T cells application (1-2 x 105 cells/well). Rotational and translational offset between both imaging sessions was corrected using 3D rigid-body registration in SimpleITK.

## Statistical analysis

To assess the joint effect of assistance on performance and task duration (**Fig. 1c**), we conducted a one-sample Hotelling’s T² test on the per-task differences (Assisted - Unassisted) in workflow agreement and completion time. To compare killing efficiency between engager states (**Fig. 2g**), a linear mixed-effects model was fitted with behavioral state as a fixed effect and random intercepts for each imaging well, each track within a well, and each track-state combination (all timepoints of a track in the same state). Then, a one way ANOVA was applied to the mixed-linear model with Satterthwaite degrees of freedom, followed by Tukey-adjusted pairwise comparisons. To compare the proportion of long-duration interactions with T cells between mesenchymal and ameboid macrophage phenotypes (**Fig. 2k**), a two-sided paired Student’s t-test was performed, with pairs defined as the ameboid and mesenchymal macrophage populations present in the same well. Behavioral state proportions of uninfected and HIV-infected T cells were compared within each cluster using a two-sided paired Student’s t-test (Extended Data Fig. 4d).

## Data availability

Data newly generated for this study will be deposited in a public repository upon publication. Previously published data reused in this study are available from their original sources indicated in Supplementary Table 2. BEHAV3D-Explorer config files containing all parameters for processing and analysis for each study shown in this work, Cellpose models, and a demo dataset are available at https://doi.org/10.5281/zenodo.18872977.

## Code availability

BEHAV3D-Explorer is released under the Apache License 2.0 at https://github.com/imAIgene-Dream3D/BEHAV3D-Explorer with documentation at https://imaigene-dream3d.github.io/BEHAV3D-Explorer/. A demo version of BEHAV3D-Explorer is available in Google Colab at https://colab.research.google.com/github/imAIgene-Dream3D/BEHAV3D-Explorer/blob/main/demo/colab/BEHAV3D_Explorer_Colab.ipynb, allowing testing without installation, though runtime is not representative of local performance.

## Acknowledgements

We acknowledge participants in the QueenB benchmarking for their time and dedication to this project. Immunotherapy co-culture imaging was performed at the Princess Máxima Imaging Center. This work was financially supported by PID2023-146455OA-I00 funded by MICIU/AEI/10.13039/501100011033 and by ERDF/EU; The Brain Tumour Charity and the WorldWide Cancer Research (2025_0170), and by the Netherlands Organization for Scientific Research (NWO) Gravitation programme IMAGINE! (project number 24.005.009 to S.B.). M.A was supported by the Comunidad de Madrid (2022-T1/BMD-24021). Development of the BEHAV3D Explorer platform and manuscript preparation was assisted by AI models, including ChatGPT, Claude, Github Copilot, Cursor and Google Gemini. All AI-generated contributions have been reviewed and edited by the authors.

## Contributions

S.d.B., E.R.J., A.R.M. and M.A. conceived the project. S.d.B., E.R.J., A.R.M. and M.A. developed the code for the BEHAV3D-Explorer platform, with contributions from O.H. and M.M.M.. S.d.B., E.R.J., A.R.M. and M.A. performed analysis of the datasets shown in this study. B.M.T.V., A.K.L.W., J.K.M. and H.F.C. performed wet-lab experiments that generated the data used in this study. S.d.B., E.R.J., A.R.M. and M.A. wrote the manuscript with revisions from all other authors. A.C.R. and M.A. supervised the study.

