## Supplementary Figures, Notes and Tables for "BEHAV3D-Explorer enables natural language-assisted profiling of 3D cellular dynamics"

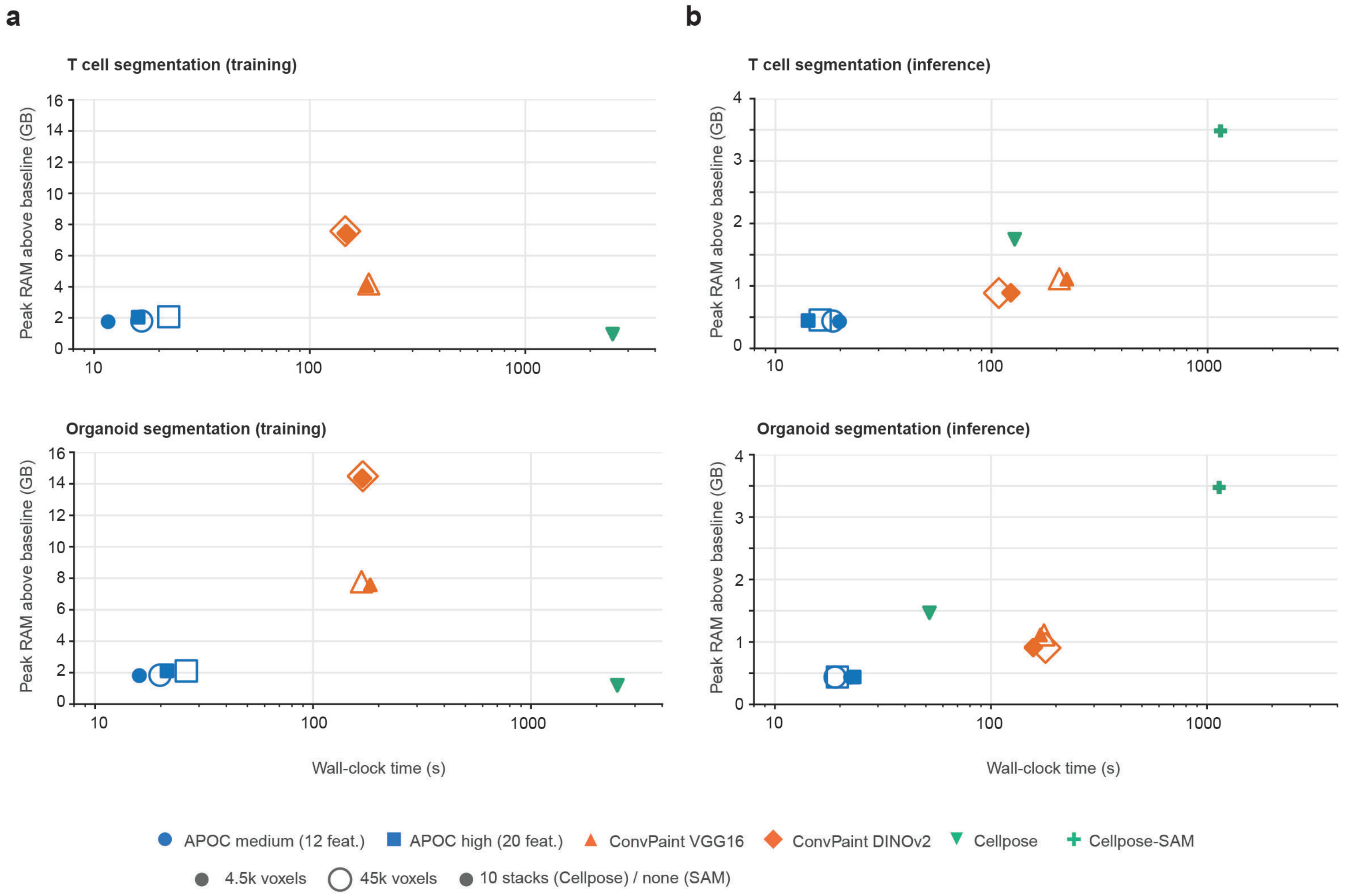


Extended Data Fig 1.

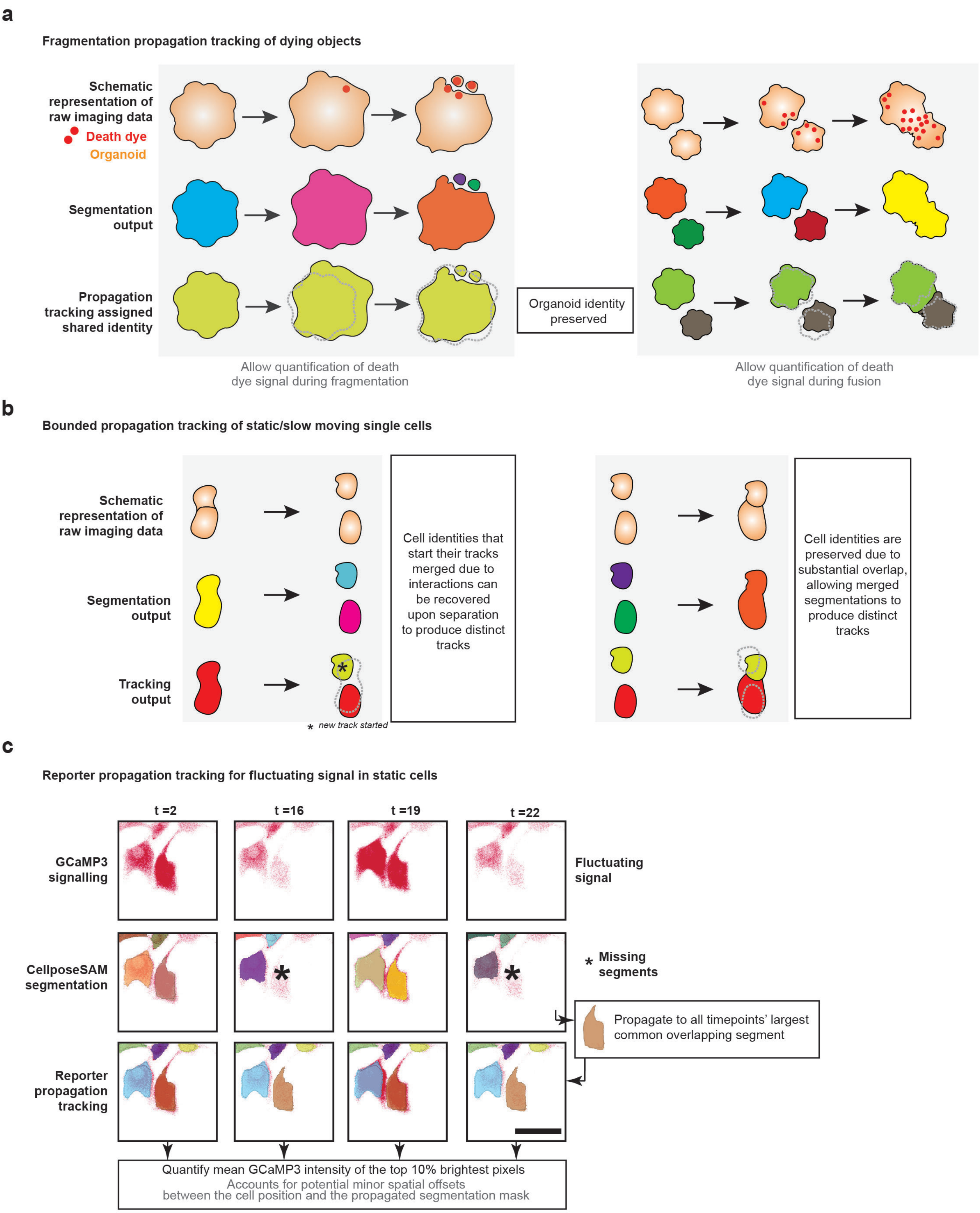


Extended Data Figure 2.

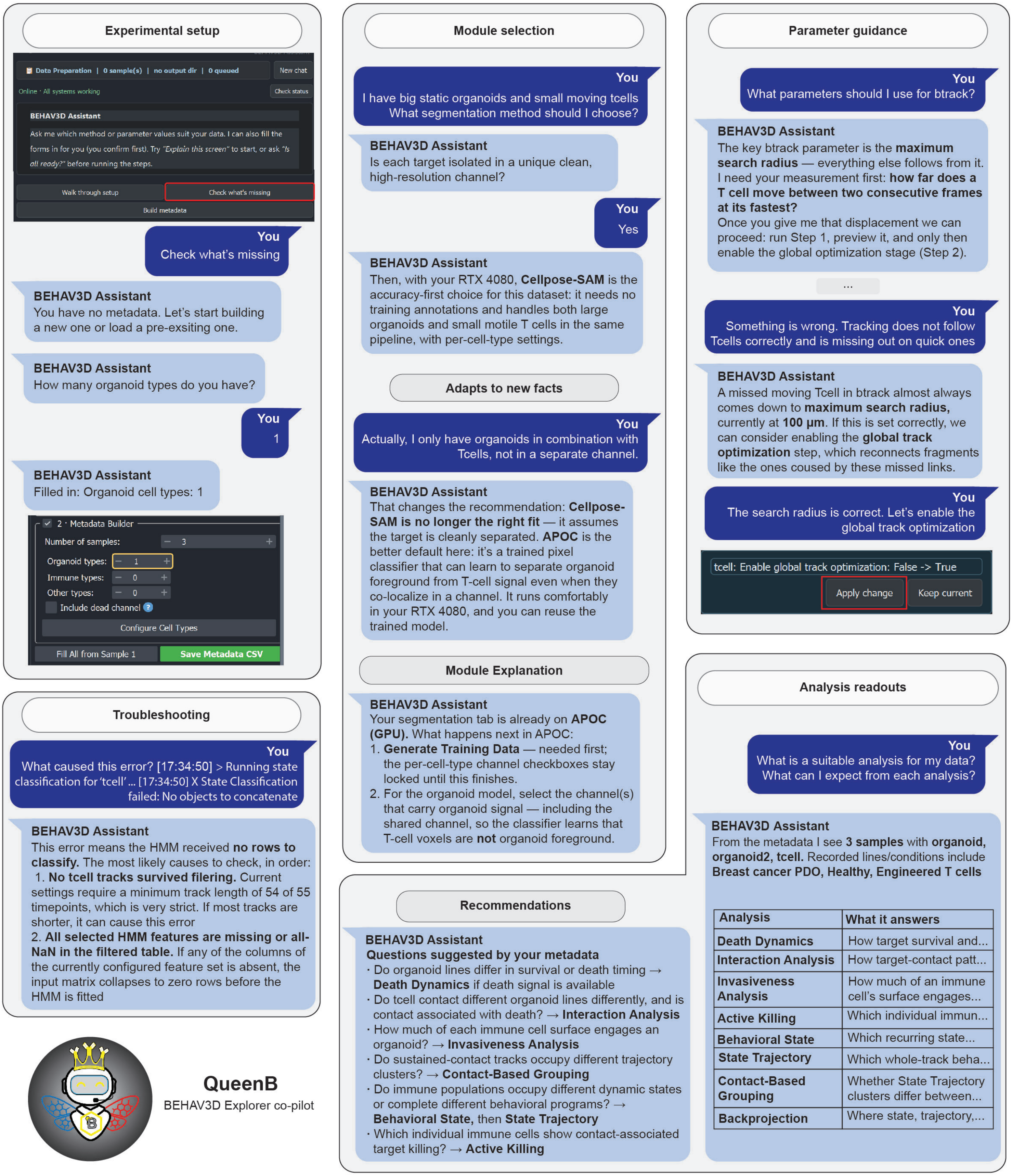


Extended Data Figure 3.

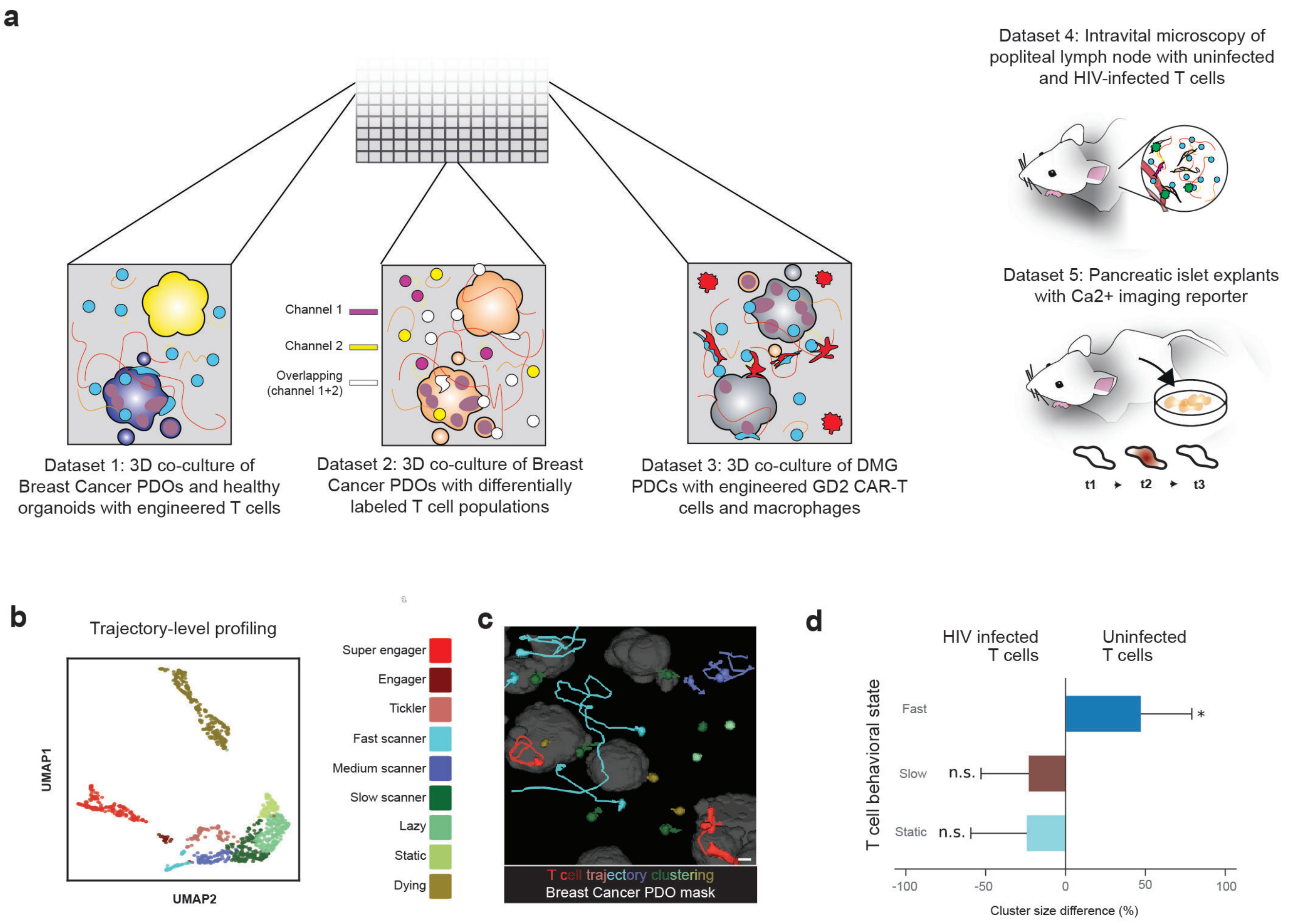


Extended Data Figure 4.

| | Reference (DOI) | This paper | 10.1038/s41467-025-57193-y | 10.7554/eLife.105302 | 10.1186/s12915-022-01372-6 | 10.1242/jcs.241422 | 10.1093/bioinformatics/btaa1106 |
|---|---|---|---|---|---|---|---|
| | **Feature** | **BEHAV3D-Explorer** | **Cecelia** | **CellDetective** | **Cell-ACDC** | **Cell Tracking Profiler** | **CellTracker** |
| **Accessibility & Adoption** | Standalone no-code installation | ☑ | ✖ | ✖ | ⚠ * | ✖ | ✖ |
| | Natural-language analysis assistant | ☑ | ✖ | ✖ | ✖ | ✖ | ✖ |
| | End-to-end analysis pipeline (raw data to final results) | ☑ | ☑ | ☑ | ☑ | ✖ | ☑ |
| | Interactive, context-aware parameter explanations | ☑ | ✖ | ✖ | ✖ | ✖ | ✖ |
| | Interactive plots | ✖ | ☑ | ✖ | ⚠ ** | ✖ | ✖ |
| | Editable AI-generated workflow configuration | ☑ | ✖ | ✖ | ✖ | ✖ | ✖ |
| | Sparse-annotation-based segmentation adaptation | ☑ | ✖ | ✖ | ✖ | ✖ | ✖ |
| | Unified analysis coding environment | ☑ | ⚠ *** | ☑ | ☑ | ✖ | ☑ |
| **Image Processing** | 3D analysis | ☑ | ☑ | ✖ | ⚠ **** | ☑ | ✖ |
| | Includes zero-shot segmentation | ☑ | ✖ | ✖ | ☑ | ✖ | ✖ |
| | Offers simultaneous multi-organoid tracking | ☑ | ✖ | ✖ | ✖ | ✖ | ✖ |
| | Handles fluctuating signal tracking (reporter fluctuation) | ☑ | ✖ | ✖ | ✖ | ✖ | ✖ |
| | Handles 3D structure fragmentation during tracking | ☑ | ✖ | ✖ | ✖ | ✖ | ✖ |
| | Supports external segmentation and tracking inputs | ☑ | ✖ | ☑ | ✖ | ✖ | ✖ |
| | Supports user manual tracking correction | ☑ | ✖ | ☑ | ☑ | ☑ | ☑ |

| | |
|---|---|
| ☑ | Yes |
| ✖ | No |
| ⚠ | Unclear; partial; or feature that required extensive human-in-the-loop involvement (see notes) |

* Standalone installation only for Windows OS
** Only in Jupyter Notebooks (requires coding)
*** Has distinct analysis and visualization environments
**** Backend-dependent (e.g. YeaZ is only 2D)

**Supplementary Table 1.** Feature comparison of BEHAV3D-Explorer against contemporary live-imaging analysis software.

| Dataset | | Image resolution | Origin | Source / repository accession | Used in |
|---|---|---|---|---|---|
| 1 | 3D co-culture of Breast Cancer PDOs and healthy organoids with engineered T cells | XY=1.77 µm<br>Z=1.77 µm<br>T=120 s<br>C=4 | Newly generated | This study; see methods | Fig. 2a-b, Supplementary Video 2 |
| 2 | 3D co-culture of Breast Cancer PDOs with differentially labeled T cell populations | XY=1.38 µm<br>Z=1.38 µm<br>T= 120 s<br>C=4 | Newly generated | This study; see methods | Fig. 2c-h, Supplementary Video 3, Extended Data Figure 4b,c |
| 3 | 3D co-culture of DMG PDCs with engineered GD2 CAR-T cells and macrophages | XY=1.77 µm<br>Z=1.77 µm<br>T=131 s<br>C=4 | Newly generated | This study; see methods | Fig. 2i-k, Supplementary Video 4 |
| 4 | Intravital microscopy of popliteal lymph node with uninfected and HIV-infected T cells | XY=1.15 µm<br>Z=4 µm<br>T=15 s<br>C=4 | Reused (published) | Murooka et al., Nature 490, 283–287 (2012); raw data via Leukocyte Tracking Database, Pizzagalli et al., Sci. Data 5, 180129 (2018) [CS002, CS003, CS006] | Extended Data Figure 4d, Supplementary Video 5 |
| 5 | Pancreatic islet explants with Ca2+ imaging reporter | XY=0.33 µm<br>Z=2 µm<br>T=5 s<br>C=2 | Newly generated | This study; see methods | Fig 2l-n, Supplementary Video 6, Extended Data Figure 2c |

**Supplementary Table 2.** Details of the five case-study datasets used to showcase BEHAV3D-Explorer analysis capabilities. Datasets marked ‘Reused (published)’ originate from external sources and are cited and linked to their original repository. Datasets marked ‘Newly generated’ were acquired for this study and are explained in the methods section. For a schematic view, see Extended Data Figure 4a.

**Supplementary Note 1.** Explanation of segmentation backends and their benchmarking.

BEHAV3D-Explorer implements segmentation and tracking as modular, interchangeable backends. This note explains that design and how the segmentation backends were benchmarked.

**Modular segmentation options and associated trade-offs**

Given that no single segmentation method performs reliably across all imaging contexts, BEHAV3D-Explorer provides multiple segmentation routes with different trade-offs in speed, annotation effort, computational costs and accuracy, benchmarked on a representative 3D live-imaging dataset (10 timepoints, 4 channels, 60 z-slices of 400 × 400 pixels, with a voxel size of 1.77 × 1.77 × 1.77 µm and a frame interval of 2 min). These routes include Cellpose-SAM[1] as a zero-shot option requiring no manual annotation, alongside several complementary, annotation-based workflows for datasets where zero-shot performance is insufficient. A key design principle of BEHAV3D-Explorer is rapid dataset-specific adaptation from sparse user annotations, allowing users to account for experiment-to-experiment variability in signal intensity, background, object density, cell size and channel bleed-through. The platform supports this through complementary sparse-annotation workflows. GPU-accelerated pixel-classification models (APOC) provide a fast route for interactive training and rapid parameter refinement on standard laboratory hardware. ConvPaint-based workflows[2] extend this strategy by combining sparse annotations with pretrained neural-network feature extractors, including vision-transformer-based DINOv2 representations, which provide broader contextual information, useful for segmenting more complex tissue-like architectures. For challenging datasets, models fine-tuned on instance annotations can improve object separation and boundary accuracy. BEHAV3D-Explorer therefore also supports trainable deep-learning segmentation based on Cellpose[3]. Finally, externally generated masks can be imported, ensuring compatibility with algorithms developed outside the BEHAV3D-Explorer platform. This modular architecture future-proofs the tool and keeps downstream biological analyses independent of the performance, availability or maintenance of any single segmentation method.

**Benchmarking modular segmentation backends in BEHAV3D-Explorer**

To guide users in selecting an appropriate workflow, we benchmarked the segmentation options available in BEHAV3D-Explorer on a single 10-timepoint 3D time-lapse acquisition containing one immune-cell (T cell) population and one organoid population. For each method we measured training time, inference time over the 10-timepoint stack, peak RAM and peak VRAM usage. These measurements were used to compare practical trade-offs in speed, annotation effort and computational costs across scenarios, not to identify a universally optimal segmentation method, in line with BEHAV3D-Explorer's fully modular and user-tailored analysis strategy

To compare sparse-annotation workflows under controlled conditions two annotation regimes were evaluated: 4,500 and 45,000 annotated voxels, representing low- and higher-annotation settings. For the two-cell-type condition, the budget was split between the T-cell and organoid layers proportionally.

We evaluated one-class and two-class segmentation settings. In the one-class setting, models were trained to segment a single target population (T cell or organoid). In the two-class setting, models were trained to segment both populations simultaneously. Because some workflows train all classes jointly whereas others handle one class at a time, the benchmark was structured to support practical comparison across implementation styles: ConvPaint trains a single unified multiclass model, so its two-class condition fits one joint model spanning both populations, while APOC fits one binary classifier per declared cell type, meaning its two-class condition trains two independent classifiers. One class and two-class segmentations were evaluated with low and high annotation settings.

We compared feature tiers reflecting realistic BEHAV3D-Explorer use. For APOC sparse pixel classification, we evaluated medium- and high-feature configurations (12 and 20 features respectively, both fit with a 100-tree random forest). For ConvPaint-based workflows, we evaluated a VGG16 feature extractor (default) and a DINOv2 feature extractor. VGG16-derived convolutional features capture local image properties such as edges, textures and channel relationships, while DINOv2-derived vision-transformer features provide broader contextual representations. In BEHAV3DExplorer both pixel-classifier types are converted from class probabilities to instance masks by the same strategy, so post-processing will not contribute to the measured differences between them.

Cellpose was evaluated separately as a trainable deep-learning instance-segmentation workflow, fine-tuned per cell type for 400 epochs from a matched pretrained backbone (*cyto3* for organoids, *nuclei* for T cells) on 10 fully annotated 3D stacks (300 annotated 2D planes), then applied to the same 10 timepoints as the other methods. This comparison was included to estimate the practical computational cost of using a trainable instance-segmentation model when users require improved object separation or boundary definition compared to the

sparse-annotation workflows. However, the substantially higher annotation effort required for Cellpose was not considered in this analysis, as annotation time is highly dependent on the user.
Cellpose-SAM (pretrained cpsam) was applied zero-shot at default settings and required no annotation or training step.
All benchmarks were run on a single workstation (AMD Ryzen Threadripper PRO 5955WX, 32 logical cores; 128 GB RAM; NVIDIA RTX 4000 SFF Ada Generation, 20 GB; Windows 10). Runs were executed one at a time and timepoints were processed sequentially, so reported runtimes reflect each method alone and without competition from other applications for CPU, GPU or memory. Each condition (method × configuration × target × annotation budget) was repeated three times for training and three times for inference.

**Results**
Benchmark results across all conditions are reported below, as mean ± SD over three repeats. Inference metrics correspond to the 10-timepoint benchmark stack (400 × 400 × 60 voxels), and peak RAM and VRAM usage are reported separately for training and inference. Dashes indicate cases where no training phase was required (zero-shot Cellpose-SAM).

| Target | Method | Feature tier | Classes | Annotation | Train (s) | Inference (s) | RAM train (GB) | RAM infer. (GB) | VRAM train (GB) | VRAM infer. (GB) |
|---|---|---|---|---|---|---|---|---|---|---|
| T cell | APOC | medium (12 feat.) | 1x1 (bg+fg) | 4 500 voxels | 11.60 ± 0.07 | 19.79 ± 7.82 | 1.757 ± 0.002 | 0.429 ± 0.009 | 0.893 ± 0.000 | 0.445 ± 0.000 |
| T cell | APOC | medium (12 feat.) | 1x1 (bg+fg) | 45 000 voxels | 16.62 ± 0.04 | 18.45 ± 6.37 | 1.802 ± 0.013 | 0.437 ± 0.011 | 0.893 ± 0.000 | 0.447 ± 0.000 |
| T cell | APOC | high (20 feat.) | 1x1 (bg+fg) | 4 500 voxels | 16.00 ± 0.10 | 14.16 ± 1.29 | 2.047 ± 0.006 | 0.445 ± 0.004 | 1.486 ± 0.000 | 0.742 ± 0.000 |
| T cell | APOC | high (20 feat.) | 1x1 (bg+fg) | 45 000 voxels | 22.18 ± 0.05 | 16.12 ± 2.95 | 2.079 ± 0.008 | 0.454 ± 0.019 | 1.486 ± 0.000 | 0.744 ± 0.000 |
| T cell | ConvPaint | VGG16 | 1 (bg+fg) | 4 500 voxels | 184.30 ± 8.78 | 223.11 ± 35.72 | 4.075 ± 0.001 | 1.108 ± 0.005 | 0.566 ± 0.000 | 0.566 ± 0.000 |
| T cell | ConvPaint | VGG16 | 1 (bg+fg) | 45 000 voxels | 187.83 ± 6.25 | 206.12 ± 70.21 | 4.182 ± 0.036 | 1.115 ± 0.004 | 0.566 ± 0.000 | 0.566 ± 0.000 |
| T cell | ConvPaint | DINOv2 | 1 (bg+fg) | 4 500 voxels | 147.39 ± 0.09 | 122.85 ± 34.55 | 7.429 ± 0.003 | 0.890 ± 0.017 | 0.107 ± 0.000 | 0.107 ± 0.000 |
| T cell | ConvPaint | DINOv2 | 1 (bg+fg) | 45 000 voxels | 146.09 ± 1.29 | 108.15 ± 23.94 | 7.572 ± 0.072 | 0.886 ± 0.024 | 0.107 ± 0.000 | 0.107 ± 0.000 |
| T cell | Cellpose | not applicable | instance (no classes) | 10 annotated stacks | 2541.21 ± 82.74 | 128.10 ± 0.26 | 0.959 ± 0.001 | 1.746 ± 0.003 | 2.132 ± 0.000 | 0.515 ± 0.000 |
| T cell | Cellpose-SAM | pretrained cpsam | instance (no classes) | none (zero-shot) | – | 1148.87 ± 1.60 | – | 3.484 ± 0.003 | – | 1.718 ± 0.000 |
| Organoid | APOC | medium (12 feat.) | 1x1 (bg+fg) | 4 500 voxels | 15.96 ± 0.03 | 22.40 ± 0.95 | 1.814 ± 0.003 | 0.435 ± 0.008 | 0.888 ± 0.009 | 0.447 ± 0.000 |
| Organoid | APOC | medium (12 feat.) | 1x1 (bg+fg) | 45 000 voxels | 19.85 ± 0.12 | 18.99 ± 1.69 | 1.843 ± 0.017 | 0.437 ± 0.001 | 0.893 ± 0.000 | 0.447 ± 0.000 |
| Organoid | APOC | high (20 feat.) | 1x1 (bg+fg) | 4 500 voxels | 21.39 ± 0.18 | 23.29 ± 0.09 | 2.118 ± 0.021 | 0.439 ± 0.007 | 1.486 ± 0.000 | 0.744 ± 0.000 |
| Organoid | APOC | high (20 feat.) | 1x1 (bg+fg) | 45 000 voxels | 26.21 ± 0.11 | 19.46 ± 1.31 | 2.112 ± 0.005 | 0.439 ± 0.002 | 1.486 ± 0.001 | 0.744 ± 0.000 |
| Organoid | ConvPaint | VGG16 | 1 (bg+fg) | 4 500 voxels | 182.98 ± 4.85 | 169.51 ± 5.42 | 7.579 ± 0.021 | 1.112 ± 0.001 | 0.566 ± 0.000 | 0.566 ± 0.000 |
| Organoid | ConvPaint | VGG16 | 1 (bg+fg) | 45 000 voxels | 166.85 ± 1.03 | 175.92 ± 24.55 | 7.778 ± 0.067 | 1.112 ± 0.002 | 0.566 ± 0.000 | 0.566 ± 0.000 |
| Organoid | ConvPaint | DINOv2 | 1 (bg+fg) | 4 500 voxels | 168.48 ± 0.05 | 156.94 ± 31.88 | 14.351 ± 0.116 | 0.911 ± 0.006 | 0.107 ± 0.000 | 0.107 ± 0.000 |
| Organoid | ConvPaint | DINOv2 | 1 (bg+fg) | 45 000 voxels | 169.05 ± 1.12 | 178.83 ± 45.58 | 14.481 ± 0.027 | 0.905 ± 0.012 | 0.107 ± 0.000 | 0.107 ± 0.000 |
| Organoid | Cellpose | not applicable | instance (no classes) | 10 annotated stacks | 2489.81 ± 3.52 | 51.92 ± 0.05 | 1.195 ± 0.428 | 1.466 ± 0.006 | 2.081 ± 0.001 | 0.515 ± 0.000 |
| Organoid | Cellpose-SAM | pretrained cpsam | instance (no classes) | none (zero-shot) | – | 1138.46 ± 1.84 | – | 3.476 ± 0.011 | – | 1.718 ± 0.000 |
| T cell + Organoid | APOC | medium (12 feat.) | 2x1 (bg+fg) | 4 500 voxels | 26.94 ± 0.18 | 59.03 ± 6.85 | 2.476 ± 0.076 | 0.441 ± 0.011 | 0.893 ± 0.000 | 0.447 ± 0.000 |
| T cell + Organoid | APOC | medium (12 feat.) | 2x1 (bg+fg) | 45 000 voxels | 31.89 ± 0.07 | 61.60 ± 1.38 | 2.452 ± 0.026 | 0.443 ± 0.006 | 0.888 ± 0.009 | 0.447 ± 0.000 |

| Target | Method | Feature tier | Classes | Annotation | Train (s) | Inference (s) | RAM train (GB) | RAM infer. (GB) | VRAM train (GB) | VRAM infer. (GB) |
|---|---|---|---|---|---|---|---|---|---|---|
| T cell + Organoid | APOC | high (20 feat.) | 2x1 (bg+fg) | 4 500 voxels | 36.31 ± 0.16 | 42.47 ± 4.12 | 2.729 ± 0.073 | 0.454 ± 0.009 | 1.486 ± 0.001 | 0.744 ± 0.000 |
| T cell + Organoid | APOC | high (20 feat.) | 2x1 (bg+fg) | 45 000 voxels | 42.59 ± 0.11 | 56.58 ± 7.03 | 2.709 ± 0.005 | 0.464 ± 0.012 | 1.486 ± 0.000 | 0.744 ± 0.000 |
| T cell + Organoid | ConvPaint | VGG16 | 2 (bg+fg1+fg2) | 4 500 voxels | 334.05 ± 3.68 | 243.75 ± 55.03 | 20.484 ± 0.148 | 1.402 ± 0.007 | 0.616 ± 0.000 | 0.616 ± 0.000 |
| T cell + Organoid | ConvPaint | VGG16 | 2 (bg+fg1+fg2) | 45 000 voxels | 295.43 ± 3.51 | 288.38 ± 70.46 | 20.728 ± 0.123 | 1.365 ± 0.006 | 0.616 ± 0.000 | 0.616 ± 0.000 |
| T cell + Organoid | ConvPaint | DINOv2 | 2 (bg+fg1+fg2) | 4 500 voxels | 298.43 ± 1.69 | 346.28 ± 60.37 | 39.337 ± 0.058 | 0.981 ± 0.006 | 0.107 ± 0.000 | 0.107 ± 0.000 |
| T cell + Organoid | ConvPaint | DINOv2 | 2 (bg+fg1+fg2) | 45 000 voxels | 299.24 ± 1.86 | 221.02 ± 8.52 | 39.730 ± 0.179 | 0.872 ± 0.205 | 0.107 ± 0.000 | 0.107 ± 0.000 |

**Supplementary Note 2.** QueenB evaluation study design.

**Dataset**

The eight tasks selected for QueenB benchmarking span key decisions a new user must make to transform a raw dataset into an interpretable biological readout. All tasks were performed on the same four-channel live-imaging dataset of organoid-T cell co-cultures, containing two organoid classes (healthy and tumor, co-cultured in the same well in a subset of samples), T cells, and a dead-cell marker. Imaging was at 1.77 µm isotropic sampling with a 2 min interval across five wells. Before the first scored task, each participant received a fixed orientation covering the dataset, the layout of the interface, how to invoke QueenB, and the location of the github and wiki documentation.

**Tasks**

**T1 Understanding of the software.** "How can you use this tool? And what type of analysis can you do?" *Starting state:* fresh setup, no metadata loaded. *Success criteria:* participant has understood and can explain the steps and outputs in the software.

**T2 Set up the metadata for your experiment.** "You need to set up the metadata for your experiment so you can process it in BEHAV3D Explorer. This metadata tells BEHAV3D Explorer what samples are in your experiment, what imaging settings were used and what types of cells and organoids are in your data. Make sure to fully set up the metadata." *Starting state:* fresh setup, no metadata loaded. Channel assignments, voxel size, time interval and file names were supplied. *Success criteria:* metadata is fully filled; the two organoid classes are declared as separate types, since they share a well and require separate segmentation; the dead-cell channel is included.

**T3 Choose a segmentation method.** "You need to segment your cells before anything else. Choose which segmentation method to use." *Starting state:* metadata loaded only, no segmentation or training data. *Success criteria:* participant selects a method and can state the reasoning behind it in their own words.

**T4 Tune segmentation parameters (APOC, probability map + watershed).** "You've chosen APOC with Probability map + watershed as a segmentation method. Now tune the parameters correctly for each cell type: MDO, 27T, and T cells." *Starting state:* all parameter fields preset with intentionally incorrect defaults, so no field can be left untouched. *Success criteria:* parameters accurately set per cell type.

**T5 Choose and set up tracking method, per cell type.** "Segmentation is done. Now choose the right tracking method for healthy and tumor organoids, and for T cells, and configure it. T cells are fast moving cells. Organoids are more static structures, but some of them die and get fragmented." *Starting state:* segmentation masks completed and saved; starting in the tracking tab. *Success criteria:* correct method and parameters for each of the three cell/object types.

**T6 Configure an active-killing analysis.** “Some T cells contact an organoid without consequence, whereas after contact by others the organoid starts to die. "Configure the active killing analysis to find out if there is a difference in the rate of T cells actively killing healthy organoids versus the tumor organoids. From looking at the movie, you observe that organoids die around 30 minutes after the initial contact. You want to know if at least 1 cell dies after the T cell contacted the organoid." *Starting state:* tracking and feature extraction complete, features filtered. *Success criteria:* participant configures an analysis that quantifies active killing per target type.

**T7 Behavioral analysis of T cells.** "You want to characterize the different types of behavior your T cells show, using movement, morphology, engagement with tumor organoids and active killing, over a ~10-min window.” *Starting state:* tracking and feature extraction complete, features filtered, starting in the Analysis tab. *Success criteria:* per-timepoint behavioral classification configured with tumor organoid_ contact and active_killing as binary groups.

**T8 Understand an error.** "You ran the behavioral state analysis but you get an error in the log. Figure out what was wrong and how to fix this error." *Starting state:* tracking and feature extraction complete, features filtered wrongly, leaving no single T cell. We start in the behavioral state classification, we run it and the log gives an error. *Success criteria:* participant explains the cause of the error and fixes it.

**Limitations**

With only eight participants, individual background contributed appreciable variability, both in prior microscopy experience and in prior use of large language models. We observed a trend for participants already familiar with LLM interfaces to benefit more from QueenB than those without that experience. We do not report this comparison quantitatively, because each participant contributed different tasks to the assisted and unassisted conditions and the number of participants per experience is too small to support an objective conclusion. Time was capped at 30 min per task. The cap was reached on several occasions, all of them in the unassisted condition, so the unassisted distribution is right-truncated and the reported time reduction is conservative. All tasks used a single co-culture dataset to avoid more variability however generalization to other imaging modalities underlies QueenB's knowledge base but was not tested directly.

**Results**

Per-task and per-participant measurements of annotation time, agreement with the gold standard, and self-reported confidence, together with exclusion flags and the corresponding reasons for exclusion, are reported below.

| Task | Condition | User | Moder-ator | Duration (mm:ss) | Matches gold standard | Confidence (1-10) | Exclude | Exclusion reason |
|---|---|---|---|---|---|---|---|---|
| T1 | Assisted | P1 | M1 | 8:04 | Yes | 9 | | |
| T1 | Assisted | P3 | M1 | 4:20 | Yes | 7 | | |
| T1 | Assisted | P5 | M1 | 9:30 | Yes | 7 | | |
| T1 | Assisted | P7 | M2 | 9:24 | Partial | 8 | | |
| T1 | Unassisted | P2 | M2 | 21:30 | Partial | 6 | | |
| T1 | Unassisted | P4 | M2 | 14:28 | Partial | 6 | | |
| T1 | Unassisted | P6 | M3 | 25:30 | Partial | 8 | | |
| T1 | Unassisted | P8 | M3 | 23:03 | Yes | 9 | | |
| T2 | Assisted | P1 | M1 | - | - | - | X | Temporary chatbot unavailability |
| T2 | Assisted | P5 | M1 | 13:18 | Yes | 7 | | |
| T2 | Assisted | P6 | M3 | 11:00 | Yes | 9 | | |
| T2 | Assisted | P8 | M3 | 13:37 | Yes | 10 | | |
| T2 | Unassisted | P2 | M2 | 30:00 | No | 6 | | |
| T2 | Unassisted | P3 | M1 | 20:09 | Yes | 6 | | |
| T2 | Unassisted | P4 | M2 | 20:00 | Partial | 8 | | |
| T2 | Unassisted | P7 | M3 | 26:55 | Partial | 8 | | |
| T3 | Assisted | P1 | M1 | 6:41 | Yes | 7 | | |
| T3 | Assisted | P2 | M2 | 5:00 | Yes | 10 | | |
| T3 | Assisted | P6 | M3 | - | - | - | X | Misinterpretation of task objective |
| T3 | Assisted | P8 | M3 | 3:34 | Yes | 10 | | |
| T3 | Unassisted | P3 | M1 | 9:55 | Yes | 8 | | |
| T3 | Unassisted | P4 | M2 | 9:00 | Partial | 8 | | |
| T3 | Unassisted | P5 | M1 | 9:47 | Partial | 6 | | |
| T3 | Unassisted | P7 | M3 | 21:26 | Yes | 6 | | |
| T4 | Assisted | P1 | M1 | - | - | - | X | Temporary chatbot unavailability |
| T4 | Assisted | P2 | M2 | - | - | - | X | Misinterpretation of task objective |
| T4 | Assisted | P3 | M1 | 21:11 | Yes | 8 | | |
| T4 | Assisted | P7 | M3 | 7:26 | Yes | 8 | | |
| T4 | Assisted | P8 | M3 | 12:11 | Partial | 0 | | |

| | | | | | | | | |
|---|---|---|---|---|---|---|---|---|
| T4 | Unassisted | P4 | M2 | 23:00 | Partial | 4 | | |
| T4 | Unassisted | P5 | M1 | 24:14 | Yes | 6 | | |
| T4 | Unassisted | P6 | M3 | 24:13 | Yes | 6 | | |
| T5 | Assisted | P2 | M2 | 11:00 | Yes | 10 | | |
| T5 | Assisted | P3 | M1 | 9:11 | Yes | 8 | | |
| T5 | Assisted | P4 | M2 | - | - | - | X | Misinterpretation of task objective |
| T5 | Assisted | P7 | M3 | 15:28 | Yes | 8 | | |
| T5 | Assisted | P8 | M3 | 5:44 | Yes | 10 | | |
| T5 | Unassisted | P1 | M1 | 11:33 | Partial | 6 | | |
| T5 | Unassisted | P5 | M1 | 15:52 | Partial | 7 | | |
| T5 | Unassisted | P6 | M3 | 9:46 | Partial | 8 | | |
| T6 | Assisted | P3 | M1 | 6:44 | Yes | 9 | | |
| T6 | Assisted | P4 | M2 | 5:54 | Yes | 6 | | |
| T6 | Assisted | P5 | M1 | 9:13 | Yes | 9 | | |
| T6 | Assisted | P7 | M3 | 11:24 | Yes | 8 | | |
| T6 | Unassisted | P1 | M1 | 15:00 | Yes | 7 | | |
| T6 | Unassisted | P2 | M2 | 12:51 | Yes | 8 | | |
| T6 | Unassisted | P6 | M3 | 16:17 | Partial | 8 | | |
| T6 | Unassisted | P8 | M3 | 25:38 | Partial | 8 | | |
| T7 | Assisted | P4 | M2 | 12:54 | Yes | 8 | | |
| T7 | Assisted | P5 | M1 | 17:21 | Partial | 7 | | |
| T7 | Assisted | P6 | M3 | 15:02 | Yes | 8 | | |
| T7 | Unassisted | P1 | M1 | 19:07 | Yes | 7 | | |
| T7 | Unassisted | P2 | M2 | 30:00 | Partial | 6 | | |
| T7 | Unassisted | P3 | M1 | 23:46 | Partial | 10 | | |
| T7 | Unassisted | P7 | M3 | 23:56 | Partial | 6 | | |
| T7 | Unassisted | P8 | M3 | 19:04 | Partial | 9 | | |
| T8 | Assisted | P2 | M2 | 6:25 | Yes | 10 | | |
| T8 | Assisted | P4 | M2 | 5:30 | Yes | 8 | | |
| T8 | Assisted | P6 | M3 | 2:48 | Yes | 10 | | |
| T8 | Unassisted | P1 | M1 | 12:55 | Yes | 9 | | |
| T8 | Unassisted | P3 | M1 | - | - | - | X | Incomplete task briefing by moderator |
| T8 | Unassisted | P5 | M1 | 17:53 | Yes | 8 | | |
| T8 | Unassisted | P7 | M3 | 18:53 | Yes | 6 | | |
| T8 | Unassisted | P8 | M3 | - | - | - | X | Task not completed (participant had to leave session) |
